# Novel Insights into the Aortic Mechanical Properties of Mice Modeling Hereditary Aortic Diseases

**DOI:** 10.1101/2023.08.15.553452

**Authors:** Nicolo Dubacher, Kaori Sugiyama, Jeffrey D. Smith, Vanessa Nussbaumer, Máté Csonka, Szilamér Ferenczi, Krisztina J. Kovács, Sylvan M. Caspar, Lisa Lamberti, Janine Meienberg, Hiromi Yanagisawa, Mary B. Sheppard, Gabor Matyas

## Abstract

**OBJECTIVE:** Hereditary aortic diseases (hADs) increase the risk of aortic dissections and ruptures. Recently, we have established an objective approach to measure the rupture force of the murine aorta, thereby explaining the outcomes of clinical studies and assessing the added value of approved drugs in vascular Ehlers-Danlos syndrome (vEDS). Here, we applied our approach to six additional mouse hAD models.

**APPROACH AND RESULT:** We used two mouse models of Marfan syndrome (MFS) as well as one smooth-muscle-cell-specific knockout (SMKO) of *Efemp2* and three CRISPR/Cas9-engineered knock-in models (*Ltbp1*, *Mfap4*, and *Timp1*). One of the two MFS models was subjected to 4-week-long losartan treatment. Per mouse, three rings of the thoracic aorta were prepared, mounted on a tissue puller, and uniaxially stretched until rupture. The aortic rupture force of the SMKO and both MFS models was significantly lower compared with wild-type mice but in both MFS models higher than in mice modeling vEDS. In contrast, the *Ltbp1*, *Mfap4*, and *Timp1* knock-in models presented no impaired aortic integrity. As expected, losartan treatment reduced aneurysm formation but surprisingly had no impact on the aortic rupture force of our MFS mice.

**CONCLUSIONS:** Our read-out system can characterize the aortic biomechanical integrity of mice modeling not only vEDS but also related hADs, allowing the aortic-rupture-force- focused comparison of mouse models. Furthermore, aneurysm progression alone may not be a sufficient read-out for aortic rupture, as antihypertensive drugs reducing aortic dilatation might not strengthen the weakened aortic wall. Our results may enable identification of improved medical therapies of hADs.

**Highlights:**

- Assay to measure the aortic rupture force as read-out for the biomechanical integrity identified weakened murine thoracic aorta prior to micro- and macroscopic changes in *Fbn1^+/C1041G^* mice.
- Despite reducing aneurysm growth, losartan treatment did not have any impact on the aortic rupture force of *Fbn1^mgR/mgR^* mice modeling MFS.
- Smooth-muscle-cell-knock-out of *Efemp2* significantly impaired the rupture force of the murine ascending aorta.
- Aortic rupture force measurements could clarify the causality of novel candidate gene(s)/variant(s) in mouse models of aortic diseases.

**Graphical Abstract:** 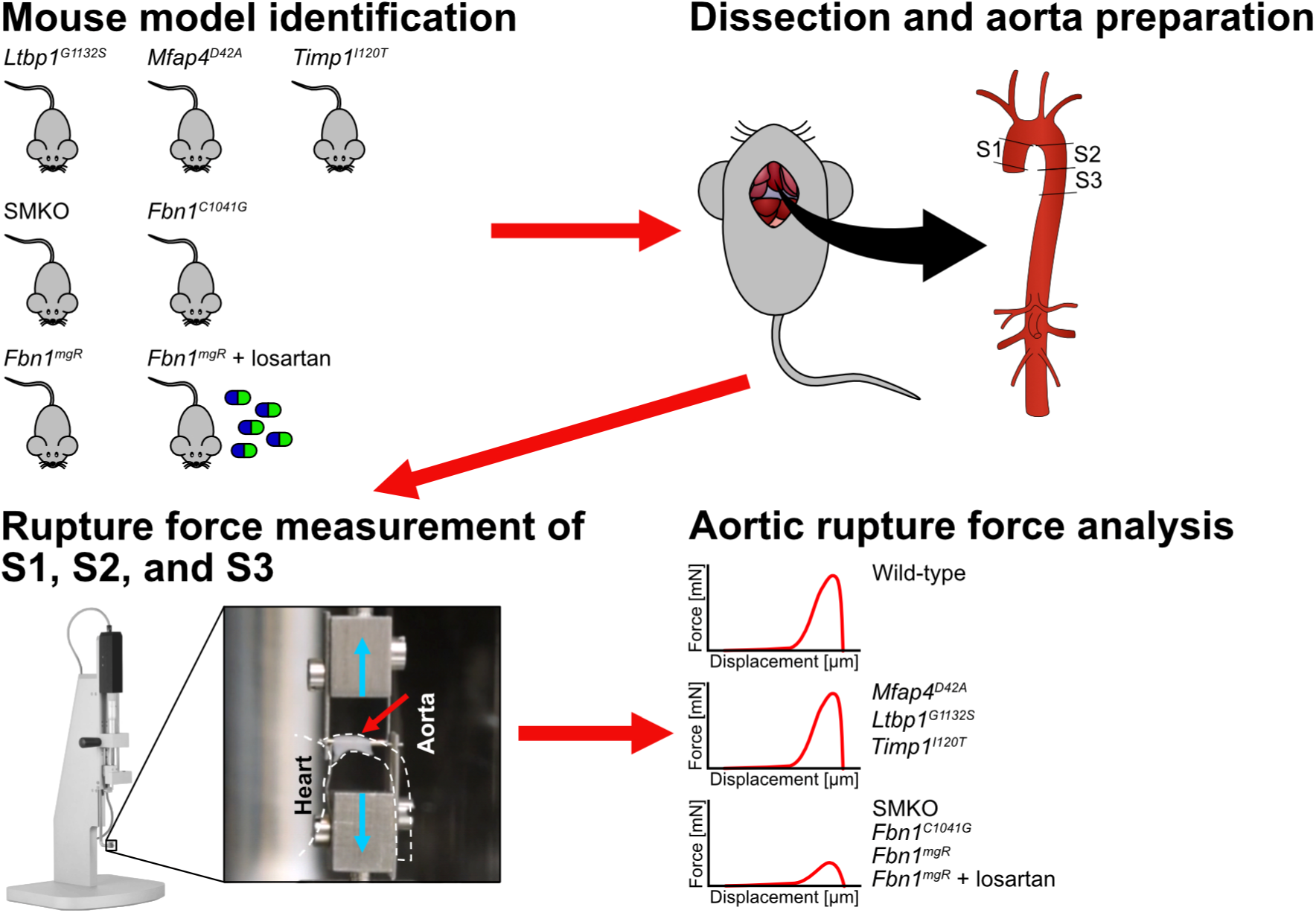

---

Hereditary aortic diseases (hADs) with life-threatening aneurysms and dissections are associated with high morbidity and mortality and are the cause for up to ∼2% of all deaths in the Western world.^1^ Syndromic and non-syndromic forms of hADs are rare (i.e., prevalence <1:2000) but several types have been identified so far. Despite considerable phenotypic similarity, disease management can profoundly differ depending on the causative gene, requiring molecular genetic testing.^2,3^ The advent of high-throughput sequencing has enabled genetic data generation at hitherto unprecedented scale, allowing for novel insights into the molecular etiology of hereditary aortopathies but requiring a high level of expertise for appropriate data analysis and variant interpretation.^4^ Currently, at least 50 genes have been associated with hereditary thoracic aortic aneurysms (TAAs) and classified based on the strength of association.^5^ However, these genes cannot explain all the cases of heritable TAAs, suggesting the presence of additional causative genes for hADs.^6^

Changes in the structural components of the extracellular matrix (ECM) as well as in chemical and mechanical stimuli can influence the molecular signaling pathways that regulate aortic wall homeostasis. Accordingly, hAD-causing sequence variants (mutations) can affect proper ECM organization, thereby directly impacting the mechanical properties of the aorta as well as the mechanoregulation of vascular smooth muscle cells and their ability to sense matrix-mediated mechanical stimuli.^7^ This can result in impaired aortic integrity and mechanical dysfunction causing progressive weakening of the aortic wall, ultimately leading to arterial dissection and/or rupture.^8^ The mechanical properties of the thoracic aorta can be derived from *in vivo* imaging parameters such as aortic wall distensibility and local pulse wave velocity or studied directly in *ex vivo* aortic samples.^9–11^ In humans, however, studies assessing the clinically relevant parameter of aortic rupture (with or without aneurysm/dissection) in hAD are significantly limited. Thus, non-human models harboring disease-causing variants in hAD-associated gene(s) provide a valuable alternative, when combined with an appropriate read-out at individual level.

Recently, we have established an objective approach to measure the aortic rupture force (maximum tensile force) of uniaxially stretched murine thoracic aortic rings as a read- out for the biomechanical integrity of the aorta.^12^ We have previously used this approach to clarify whether or not antihypertensive drugs can also strengthen the thoracic aorta (i.e., increase rupture force), thereby showing that celiprolol, a 3^rd^-generation β-blocker, but not losartan, an angiotensin-II-receptor-type-1 (AGTR1) inhibitor, can improve the biomechanical integrity of the aortic wall (i.e., beyond the celiprolol’s expected effect of slowing heart rate and lowering blood pressure).^12^ Accordingly, our approach provided a possible explanation for the previous outcomes of clinical and retrospective studies in vascular Ehlers-Danlos syndrome (vEDS, OMIM #130050).^13–15^ This added value of celiprolol, however, cannot simply be extrapolated to other β-blockers, as we have previously demonstrated for bisoprolol.^16^

Here, we applied our established read-out approach^12^ to measure the aortic rupture force in six additional mouse hAD models and compared their aortic rupture force with that of vEDS mice: (i) We used the two most widely investigated mouse models of Marfan syndrome (MFS, OMIM #154700) (*Fbn1^C1041G^* and *Fbn1^mgR^*),^17–19^ a rare autosomal-dominant inherited disease with severe cardiovascular manifestations, such as TAAs predisposing to aortic dissection and rupture, caused by mutations in the gene encoding fibrillin-1 (*FBN1* in humans, *Fbn1* in mice).^20^ In one of the two mouse MFS models (*Fbn1^mgR^*), we also aimed to clarify whether or not the frequently prescribed losartan, which has been shown to slow down aneurysm progression in mouse models as well as humans with MFS,^21–25^ also strengthens the potentially weakened thoracic aorta. (ii) Furthermore, we assessed the aortic rupture force in an *Efemp2* (encoding fibulin-4 [Fbln4]) smooth-muscle-specific knock-out (SMKO) model recapitulating key vascular phenotypic features (aortic aneurysms and arterial tortuosity)^26^ of the autosomal-recessive inherited connective tissue disorder cutis laxa type 1B (OMIM #614437).^27,28^ (iii) Moreover, we used three novel CRISPR/Cas9-engineered knock-in models (*Ltbp1^G1132S^*, *Mfap4^D42A^*, *Timp1^I120T^*) to assess the causality of human-equivalent variants in hAD candidate genes detected by whole-genome sequencing (WGS).^6^

## METHODS

### Mouse strains

In this study, multiple mouse strains were used. An overview can be found in the Major Resources Table in the Supplemental Material. Wild-type mice are referred to as *^+/+^*.

### Fbn1^C1041G^ and Fbn1^mgR^ Mouse Models

For the *Fbn1^C1041G^*mouse model (C57BL/6J background), 10-week-old heterozygous (*Fbn1^+/C1041G^*, n=10) and wild-type (*Fbn1^+/+^*, n=10) mice were ordered in frozen condition from Jackson Laboratory (JAX stock #012885, Bar Harbor, ME, USA). Tail biopsies were taken to confirm the genotypes of delivered mice, for which DNA extraction, PCR, and subsequent Sanger sequencing were performed as previously described.^12,17^

*Fbn1^mgR^* mice (C57BL/6J background) were bred and maintained on a 14/10-hour light/dark cycle receiving standard rodent chow and water *ad libitum*. Mice were genotyped as described previously.^18^ In addition, 4-week-old homozygous (*Fbn1^mgR/mgR^*, n=12) and wild-type (*Fbn1^+/+^*, n=10) mice were treated for 4 weeks with losartan dissolved in drinking water at a concentration of 600 mg/L (LTK Labs, St. Paul, MN, USA, ∼180 mg/kg bodyweight per day); age-matched, untreated *Fbn1^mgR/mgR^*(n=17) and *Fbn1^+/+^* (n=10) littermates receiving normal drinking water served as controls. Animals were weighed at the beginning and at the end of the treatment and water consumption was monitored regularly to ensure that control and treated mice drank comparable quantities. At the end of the treatment, mice were euthanized using CO_2_ and subsequently frozen. Treatment protocols were approved by the University of Kentucky IACUC (approval reference number: 2016-2437) in line with the local guidelines.

### *Efemp2* SMKO Model

*Efemp2^loxP/KO^*;*SM22-Cre^Tg/Tg^* (SMKO) and wild-type littermates (wild-type for *Efemp2* but containing the SM22α-Cre transgene, *Efemp2^+/+^*;*SM22-Cre^Tg/Tg^*) with a mixed genetic background (C57BL/6J; 129SvEv) were bred and maintained on a 12-hour light/dark cycle receiving standard rodent chow and water *ad libitum*. Genotyping was performed as previously described.^26^ Wild-type (n=11) and SMKO (n=11) littermates were euthanized with CO_2_ at 8 weeks of age and subsequently frozen. Animal experimentation and maintenance were in accordance with and approved by the Institutional Animal Experiment Committee of the University of Tsukuba, Japan (approval reference number: 21-108).

### CRISPR/Cas9-Engineered Mouse Models

Three CRISPR/Cas9 knock-in mouse models (*Ltbp1^emG1132S^*, *Mfap4^emD42A^*, and *Timp1^emI120T^*) were generated on the C57BL/6NJ background by Cyagen (Santa Clara, CA, USA), thereby introducing human-equivalent sequence variants detected in our WGS cohort of hAD patients (GRCm38/mm10 chr17(Ltbp1):g.75,315,062G>A, chr11(Mfap4):g.61,486,071A>C, and chrX(Timp1):g.20,873,395T>C, modeling human variants *LTBP1* NM_206943.4: c.3418G>A p.(Gly1140Ser), *MFAP4* NM_001198695.2: c.191A>C p.(Asp64Ala), and *TIMP1* NM_003254.3: c.356T>C p.(Ile119Thr), respectively). Founder mice of the three strains were bred and maintained on a 12-hour light/dark cycle receiving standard rodent chow and water *ad libitum*. The genotype was determined after weaning at the age of 4 weeks by PCR and subsequent Sanger sequencing using genomic DNA extracted of ear biopsies as previously described.^12^ Primers used for PCR and sequencing are listed in Table S1. One-year-old heterozygous (*Ltbp1^+/G1132S^*, n=10; *Mfap4^+/D42A^*, n=10; *Timp1^+/I120T^*, n=7), homozygous (*Ltbp1^G1132S/G1132S^*, n=10; *Mfap4^D42A/D42A^*, n=8; *Timp1^I120T/I120T^*, n=5), and hemizygous (*Timp1^0/I120T^*, n=9) mice as well as age-matched wild-type littermates serving as controls (*Ltbp1^+/+^*, n=10; *Mfap4^+/+^*, n=9; *Timp1^+/+^*, n=8) were euthanized with CO_2_ and subsequently frozen (cf. one-year-old mice were used to be able to detect any late-onset effects of the variants). Animal experimentation and maintenance were in accordance with institutional and local guidelines (approval reference number: ZH107/17, Zurich, Switzerland) conforming to the EU Directive 2010/63/EU for animal experiments.

### Aortic Rupture Force Measurements

Euthanized mice kept in a frozen state were thawed immediately before measurement of aortic rupture force. For all mouse models, the maximum tensile force at rupture (in mN) of three uniaxially stretched 1.5-mm thoracic aortic rings/segments (segment S1, ascending aorta, after heart; segment S2, descending aorta, after aortic arch; segment S3, descending aorta, adjacent to S2; cf. graphical abstract and note that S1-S3 are ring-shaped) per mouse was measured and relative rupture force was calculated by setting the arithmetic mean of the respective wild-type S1 to 100% as previously described.^12^ The stretching positions of the tissue puller until both 5 mN (load-free aortic diameter) and maximum force (rupture of aortic segment) were determined (in µm) as moved distance of the mounting pins during stretching. Mice were randomized and all measurements were performed blinded for the investigator.

### Statistical Analyses

For arithmetic means, 95% confidence intervals (CIs) were calculated using two-tailed critical values of *t*-distribution (vassarstats.net/conf_mean.html). For means with non- or slightly (<2% of means) overlapping 95% CIs compared in the text, two-tailed *P* values were calculated by two-sample *t*-test (equal sample variances according to *F*-test) or Welch’s *t*-test (unequal sample variances according to *F-*test), assuming normal distribution of independent samples (vassarstats.net/tu.html). For proportions, 95% CIs were calculated with continuity correction (vassarstats.net/prop1.html). For additional information, see Supplemental Material.

## RESULTS

### *Fbn1^C1041G^* Mouse Model

Compared to *Fbn1*^+/+^ mice, the aortic rupture force of *Fbn1^+/C1041G^*mice was significantly decreased in all three aortic segments (S1, S2, and S3) (Figure 1A; Table S2). No significant difference in stretch was detected for any of the three aortic segments of 10-week-old *Fbn1*^+/+^ and *Fbn1^+/C1041G^*mice when stretch was measured at 5 mN or maximum force (Figure 1B; Tables S3 and S4). The relative aortic rupture force and stretch values of *Fbn1*^+/+^ and *Fbn1^+/C1041G^* mice were comparable in both sexes (Figure S1).

**Figure 1.**
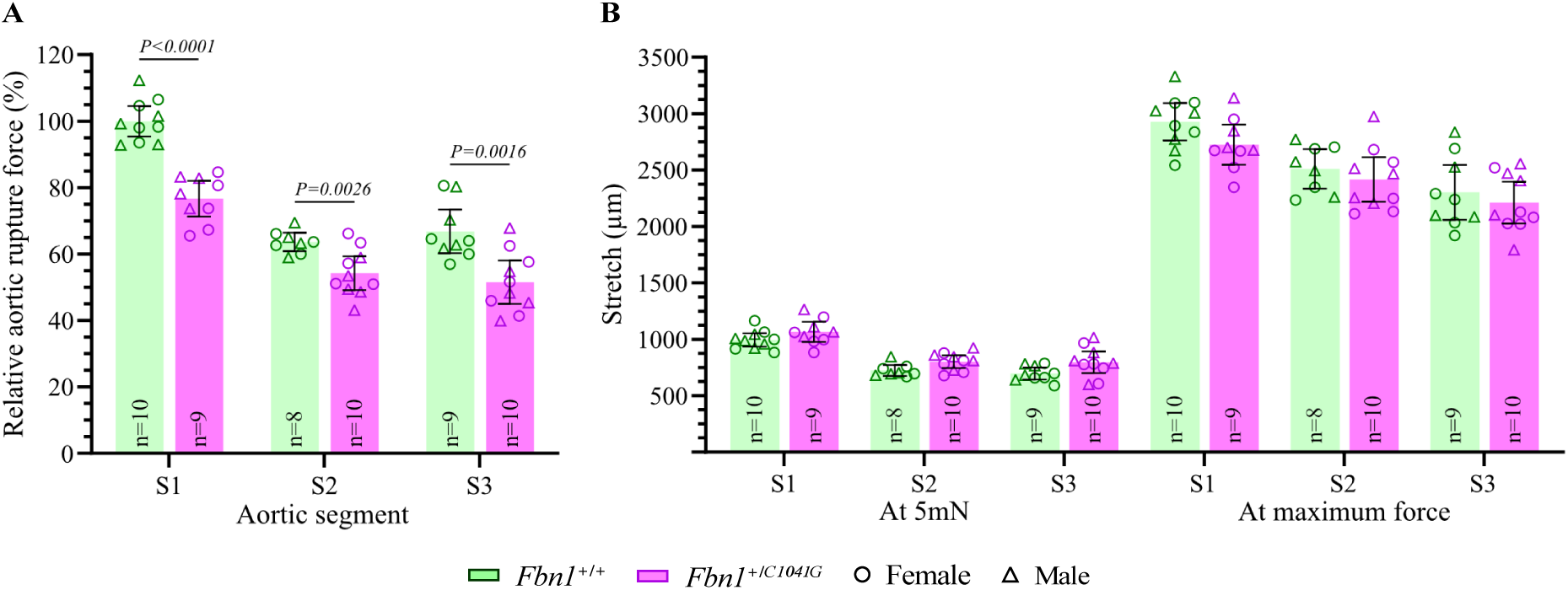
Aortic parameters of the *Fbn1^C1041G^* mouse model. A, Relative aortic rupture force (%) of three aortic segments (S1-S3) of 10-week-old *Fbn1^+/+^* and *Fbn1^+/C1041G^* mice. B, Stretch (in μm) of aortic segments at 5 mN (load-free aortic diameter) and maximum force of 10-week-old *Fbn1^+/+^* and *Fbn1^+/C1041G^* mice. Bars represent arithmetic means and error bars indicate 95% confidence intervals. The sample size (n) is displayed (damaged segments and outliers, i.e. >2× standard deviation, were excluded). For mean values with non- or slightly overlapping 95% CIs compared in text, two-tailed *P* values of unpaired t-tests are shown (applies to A only).

### *Fbn1^mgR^* Mouse Model

During 4 weeks of losartan treatment started at 4 weeks of age, the mortality rate of treated *Fbn1^mgR/mgR^* mice (8.3% [95% CI, 0.4%-40.2%]) was not significantly lower than that of untreated *Fbn1^mgR/mgR^*mice (11.8% [95% CI, 2.1%-37.7%]), but no wild-type littermate mice died (Figure 2A). On average, males were heavier than females, but body weight did not differ significantly between untreated and treated mice of the same sex and genotype either before initiation of treatment (at 4 weeks of age) or after 4 weeks of treatment (at 8 weeks of age) (Figure 2B). Of note, *Fbn1^mgR/mgR^* males were heavier than wild-type males (Figure 2B), which is unexpected considering the human MFS phenotype but has been previously observed in this mouse model and is attributed to altered adipose tissue homeostasis.^29^

**Figure 2.**
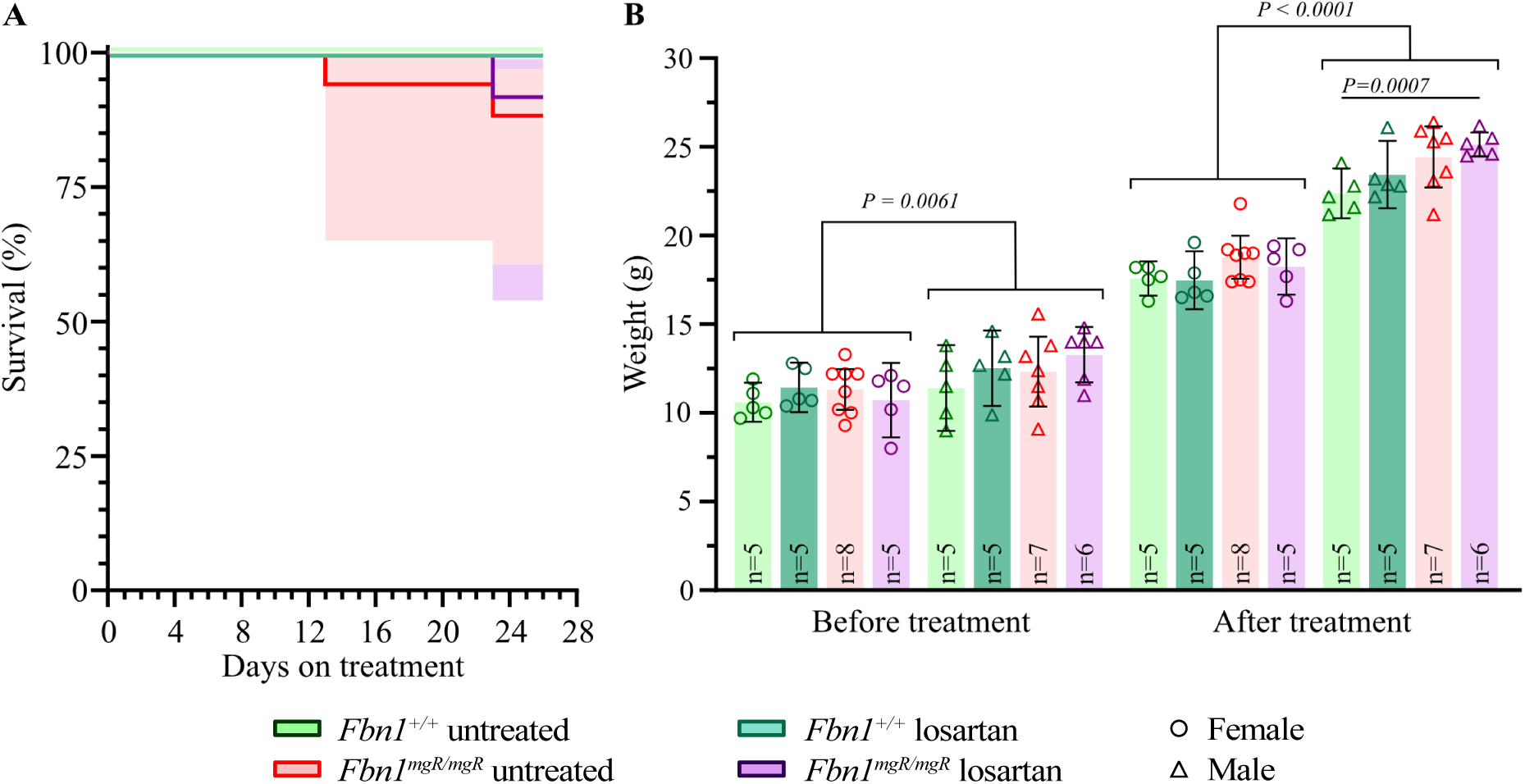
Survival and bodyweight of the *Fbn1^mgR^* mouse model treated with the AGTR1-inhibitor losartan. **A**, Kaplan-Meier curve for the survival of untreated *Fbn1^+/+^* mice (n=10) and *Fbn1^mgR/mgR^* mice (n=17), and *Fbn1^+/+^* (n=10) and *Fbn1^mgR/mgR^* (n=12) mice receiving losartan between the age of 4 and 8 weeks. Note that the survival curve of losartan-treated *Fbn1^+/+^* mice is masked by that of untreated *Fbn1^+/+^* mice. 95% confidence intervals (shaded areas) are shown for *Fbn1^mgR/mgR^* mice with survivals differing from 100%. **B**, Bodyweights of mice surviving the treatment period of 4 weeks measured before and after treatment at the age of 4 and 8 weeks, respectively. Bars represent arithmetic means, error bars indicate 95% confidence intervals. Significant two-tailed *P* values of unpaired t-tests are shown (*P*<0.05). For the comparison between female and male mice, the respective groups were pooled prior to *P* value calculation.

Compared with untreated *Fbn1^+/+^* mice, the aortic rupture force was significantly decreased in all three aortic segments of both untreated and losartan-treated *Fbn1^mgR/mgR^* mice, also revealing that losartan did not strengthen the weakened aortic wall of *Fbn1^mgR/mgR^*mice (Figure 3A; Table S5; Figure S2). The stretch of aortic segment S1 at both 5 mN and maximum force was significantly increased in untreated *Fbn1^mgR/mgR^* mice compared to untreated *Fbn1^+/+^* mice, which was more pronounced in males (Figure 3B; Tables S6 and S7). While losartan treatment of *Fbn1^+/+^* mice had no effect on S1 stretch (Figure 3B; Tables S6 and S7), losartan treatment reduced S1 stretch at both 5 mN and maximum force in *Fbn1^mgR/mgR^*mice regardless of sex (Figure 3B; Tables S6 and S7; in S2 and S3 there were no considerable differences in stretch, so these data are not shown). Overall, our data demonstrates that losartan did not significantly alter S1-S3 rupture force but did affect S1 stretch in *Fbn1^mgR/mgR^*mice modeling MFS.

**Figure 3.**
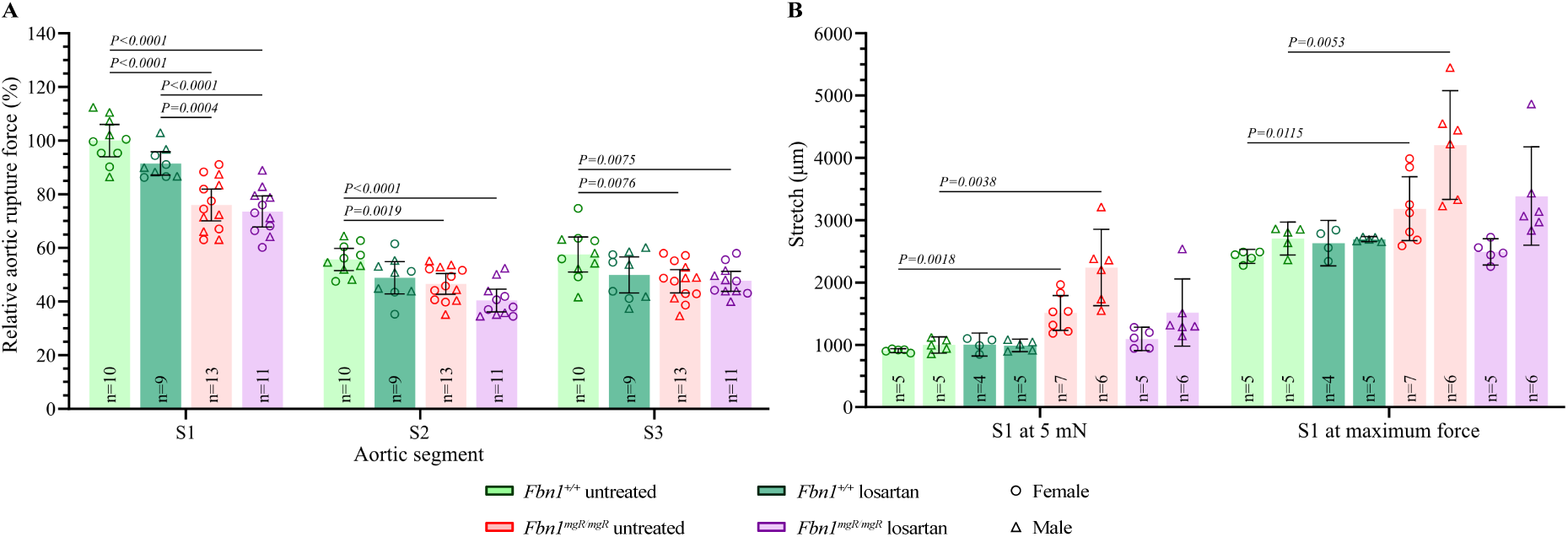
Aortic parameters of the *Fbn1^mgR^* mouse model treated with the AGTR1-inhibitor losartan. A, Relative aortic rupture force (%) of three aortic segments (S1-S3) of 8-week-old *Fbn1^+/+^* and *Fbn1^mgR/mgR^* mice after 4- week-long treatment with losartan. Untreated *Fbn1^+/+^* and *Fbn1^mgR/mgR^* mice served as control. B, Stretch (in μm) of the aortic segment S1 at 5 mN (load-free aortic diameter) as well as at maximum force in losartan-treated and untreated *Fbn1^+/+^* and *Fbn1^mgR/mgR^* mice separated according to sex. Bars represent arithmetic means and error bars indicate 95% confidence intervals (95% CI). The sample size (n) is displayed (damaged segments and outliers, i.e. >2× standard deviation, were excluded). For mean values with non- or slightly overlapping 95% CIs compared in text, two-tailed *P* values of unpaired t- tests are shown.

### *Efemp2* SMKO Mouse Model

Compared to wild-type mice, our measurements showed significantly reduced relative aortic rupture force in the ascending thoracic aorta (S1) but no difference in the descending aorta (S2 and S3) of SMKO mice (Figure 4A; Table S8). Furthermore, in SMKO mice the stretch at both 5 mN and maximum force was significantly higher in the ascending aorta (S1) but significantly lower in the descending aorta (S3) at maximum force (Figure 4B; Tables S9 and S10).

**Figure 4.**
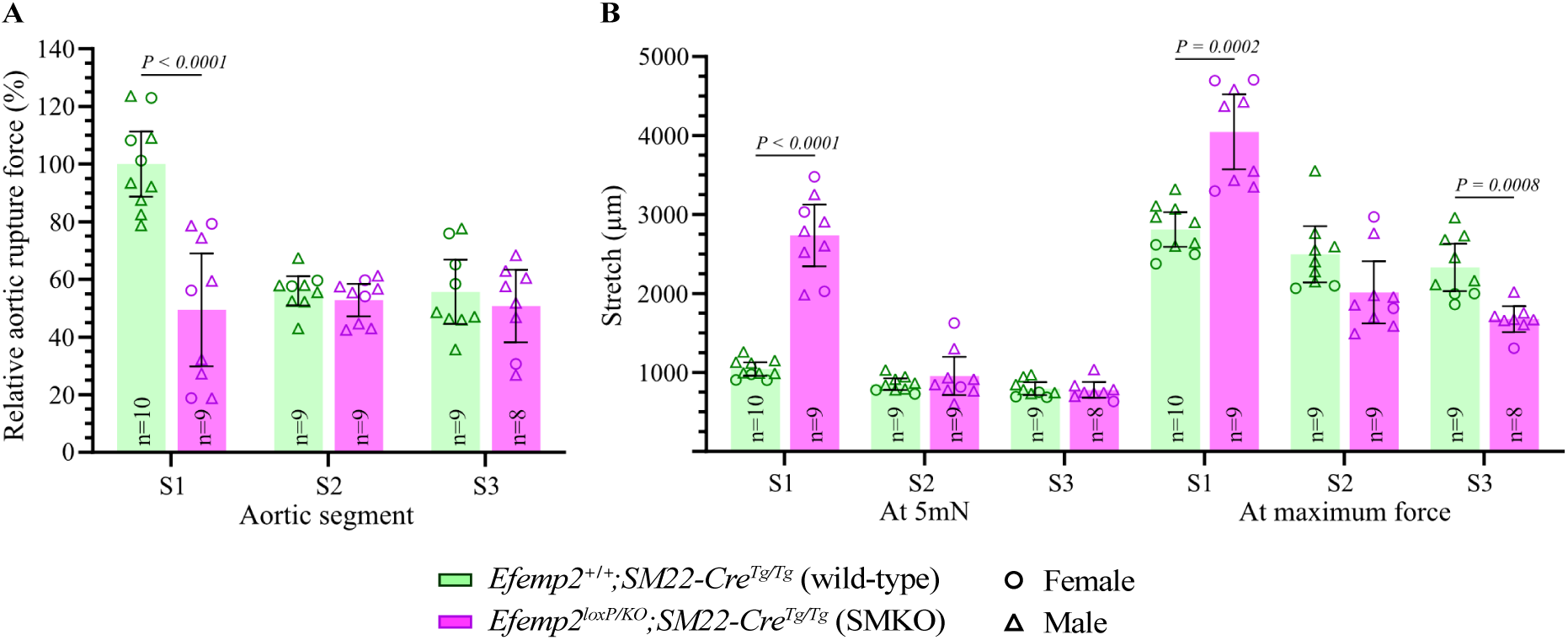
Aortic parameters of the *Efemp2* SMKO mouse model. A, Relative aortic rupture force (%) of three aortic segments (S1-S3) of 8-week-old wild-type and SMKO mice. B, Stretch (in μm) of aortic segments at 5 mN (load-free aortic diameter) as well as at maximum force in wild-type and SMKO mice. Bars represent arithmetic means and error bars indicate 95% confidence intervals. The sample size (n) is displayed (damaged segments and outliers, i.e. >2× standard deviation, were excluded). For mean values with non- or slightly overlapping 95% CIs compared in text, two-tailed *P* values of unpaired t-tests are shown.

### *Ltbp1*, *Mfap4*, and *Timp1* CRISPR/Cas9 Knock-In Mouse Models

During the one-year period of aging, no *Ltbp1*, *Mfap4*, or *Timp1* CRISPR/Cas9 knock-in mice died due to aortic rupture. Moreover, the rupture force of the aortic segments S1, S2, and S3 showed no considerable difference between wild-type controls and hetero-, homo-, or hemizygous mice of *Ltbp1*, *Mfap4*, and *Timp1* CRISPR/Cas9 knock-in mice (Figure 5; Table S11).

**Figure 5.**
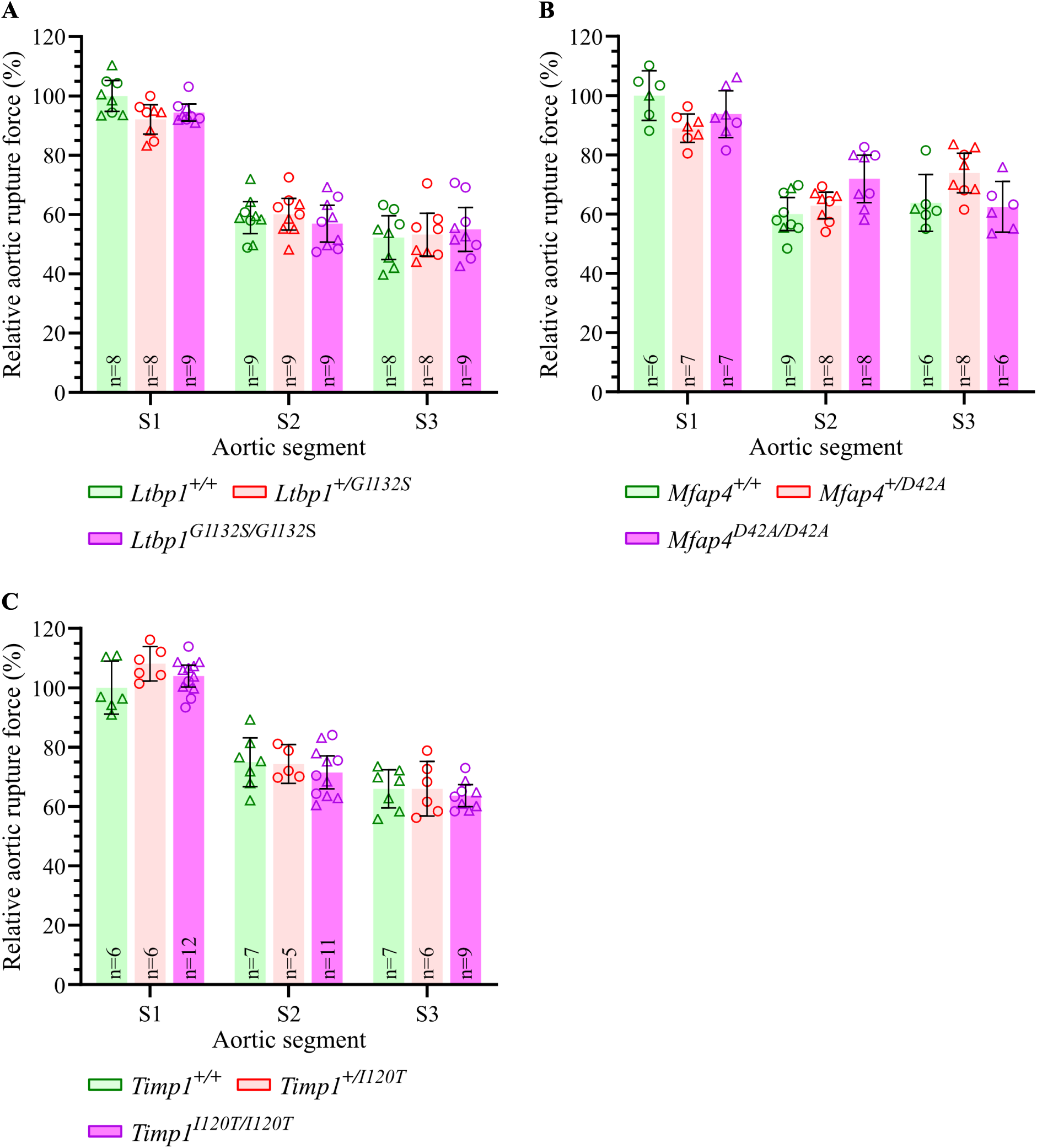
Aortic rupture force of the *Ltbp1*, *Mfap4*, and *Timp1* CRISPR/Cas9 knock-in mouse models. **A-C**, Relative aortic rupture force (%) of three aortic segments (S1-S3) of 1-year-old wild-type and hetero- as well as hemi- and homozygous *Ltbp1* (**A**), *Mfap4* (**B**), and *Timp1* (**C**) CRISPR/Cas9 knock-in mice. Bars represent arithmetic means, error bars indicate 95% confidence intervals. The sample size (n) is displayed and damaged segments and outliers (>2× standard deviation) were excluded. Note the absence of non- or slightly overlapping 95% CIs per aortic segment. Circle, female; triangle, male.

## DISCUSSION

Inbred mouse hAD models are valuable tools for the investigation of medical therapeutic approaches and the establishment of gene-disease association and have been widely adopted for the identification of cardiovascular abnormalities by assessing aortic diameter, morphology, and wall thickness as well as by considering histological staining, omics data, or survival rate.^30–36^ Although these methods provide observational insights, they cannot directly/objectively reveal how medical therapy or sequence variants influence the strength (i.e., rupture force) of the aortic wall at individual level. In this study, rupture force measurements provide insights into the biomechanical integrity of the thoracic aorta in three established (*Fbn1^C1041G^, Fbn1^mgR^*, and *Efemp2* SMKO) and three novel candidate-gene (*Ltbp1*, *Mfap4*, and *Timp1*) mouse hAD models.

By applying the aortic rupture force measurement to both mouse MFS models (i.e., the phenotypically milder heterozygous *Fbn1^C1041G^* and the phenotypically more severe homozygous *Fbn1^mgR^* mice), we showed a significantly reduced rupture force compared to wild-type littermates for the first time. The reduced rupture force is one indicator of impaired biomechanical integrity, which was most noticeably present in the ascending aorta (S1 in Figures 1A and 3A). The aortic wall of *Fbn1^+/C1041G^* mice is known to be normal in (histological) appearance around 3 months of age, only showing evidence of medial thickening and/or progressive deterioration within the medial layer, including elastic fiber fragmentation in association with excess expression of matrix metalloproteinase 2 (MMP2) and 9 (MMP9), at later ages.^17,18^ However, our read-out system allowed the identification of an impaired aortic wall integrity before 3 months of age (cf. *Fbn1^+/C1041G^*mice used in this study were 10-week-old).

Compared to wild-type controls and in contrast to *Fbn1^+/C1041G^* mice, the ascending aorta of *Fbn1^mgR/mgR^* mice showed significantly increased stretch at both 5 mN and maximum force (cf. Figure 1B and Figure 3B; note that the stretch of ring-shaped S1-S3 reflects half the circumference of the aorta and thus the aortic diameter). Increased aortic stretch was even more pronounced in male than in female mice, which is in line with previously published aortic diameters using ultrasound measurements.^37–38^ Indeed, sex is a well-known risk factor for cardiovascular diseases and several clinical studies have shown that sex also strongly influences the outcome of MFS, with men having a higher risk for aortic dilatations and arterial events.^39–41^

The antihypertensive drug losartan recognizably reduced S1 stretch, which reflects aortic diameter, likely due to lowering blood pressure (spikes), but had no impact on S1-S3 rupture force in *Fbn1^mgR/mgR^*mice (Figure 3). This result implies that the thoracic aorta treated with losartan remains susceptible to rupture despite the reduction in aortic stretch/diameter. Consequently, aortic rupture force beyond aortic diameter may be useful for the investigation of potential drug therapies in mouse models of hAD characterized by aneurysms. We propose that our read-out system provides clinically-relevant information allowing novel insights into the evaluation of medical therapies for the prevention of arterial events in hADs that cannot be obtained by assessment of the aneurysm alone.

In the *Efemp2* SMKO mouse model, our data shows that the absence of Fbln4 in the aortic ECM results in significantly reduced rupture force and increased stretch of the ascending aorta, indicating impaired biomechanical integrity of the aortic wall (Figure 4). This is likely related to the effect of Fbln4 on elastogenesis^42^ and collagen biosynthesis^26^ involved in the development of ascending aortic aneurysms,^43,44^ as previously described. Considering that the ascending aorta is subjected to the highest hemodynamic stress, it is not surprising that knock-out of *Efemp2* in vascular smooth muscle cells has a stronger effect on the ascending than on the descending aorta (i.e., S1 compared to S2 and S3), despite the higher elastin content of the ascending aorta.^45,46^

Compared to the previously reported *Col3a1^m1Lsmi^* mouse vEDS model,^12,16^ the aortic rupture forces of both *Fbn1^+/C1041G^* and *Fbn1^mgR/mgR^* mouse MFS models were significantly higher in all three aortic segments (Figure 6; Table S12). This relative aortic weakness of the *Col3a1^+/m1Lsmi^* mice is consistent with their higher mortality rate compared with *Fbn1^mgR/mgR^*mice (Figure S3; note that survival rate does not reflect aortic rupture force at the individual level, as not all mice with significantly reduced rupture force have a reduced survival). In contrast, SMKO mice showed in the ascending aorta (S1) relative aortic rupture force values comparable to or lower than those of mice modeling vEDS, but in the descending aorta (S2 and S3) the relative rupture force of SMKO mice was comparable to that of mice modeling MFS (Figure 6; Table S12).

**Figure 6.**
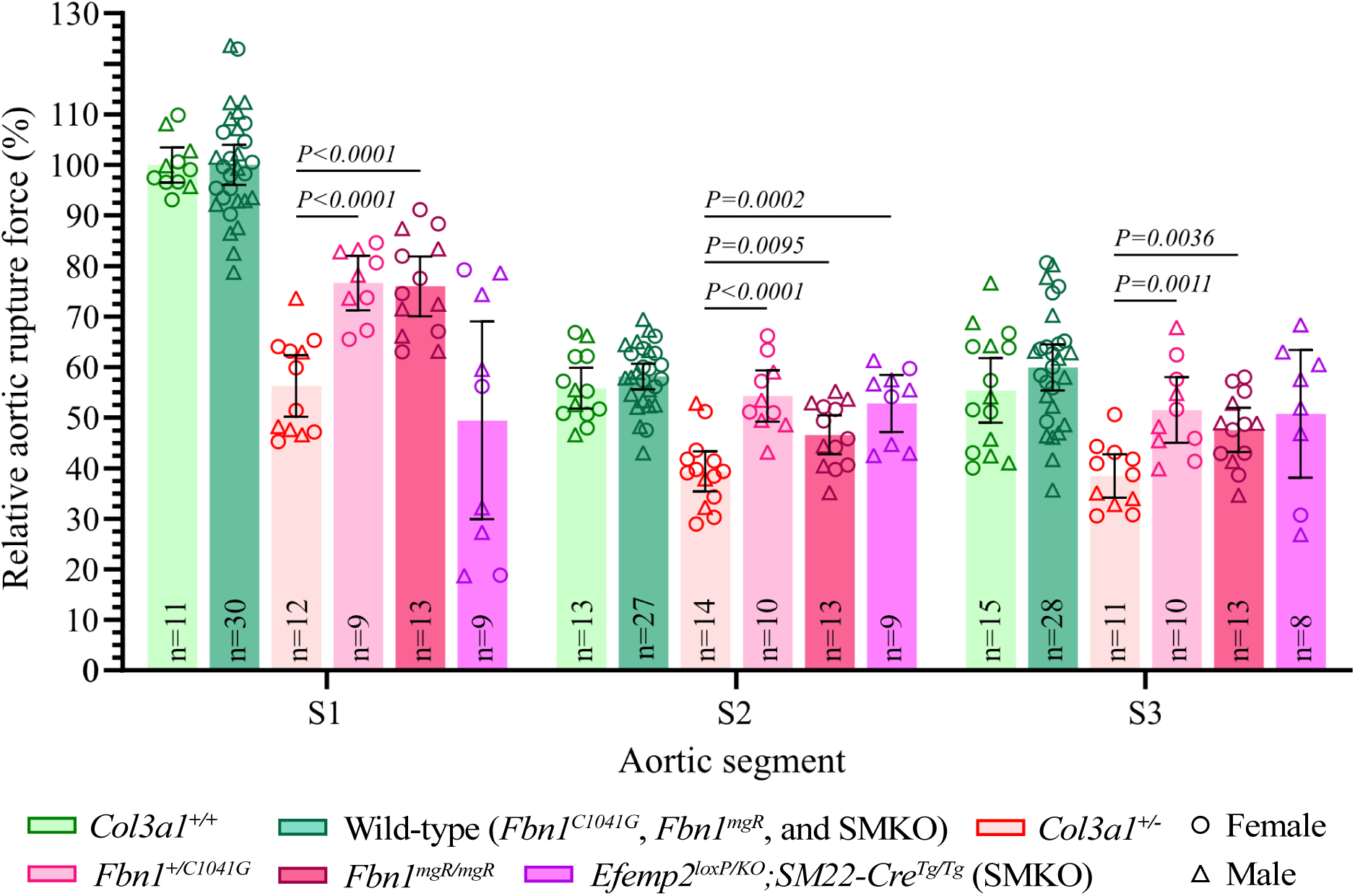
Aortic rupture force of the *Fbn1^C1041G^*, *Fbn1^mgR^*, and *Efemp2* SMKO mouse models compared to *Col3a1^m1Lsmi^* mice modeling vEDS. Relative aortic rupture force (%) of three aortic segments (S1-S3) of the mouse models *Col3a1^m1Lsmi^* (data as described previously),^12^ *Fbn1^C1041G^*, *Fbn1^mgR^*, and *Efemp2* SMKO (cf. wild-type controls of the *Fbn1^C1041G^*, *Fbn1^mgR^*, and *Efemp2* SMKO mouse models were pooled). Bars represent arithmetic means, error bars indicate 95% confidence intervals, The sample size (n) is displayed (damaged segments and outliers, i.e. >2× standard deviation, were excluded). For mean values with non- or slightly overlapping 95% CIs compared in text, two-tailed *P* values of unpaired t-tests are shown. *Col3a1^+/+^* and *Col3a1^+/-^* denotes wild-type and heterozygous mice of the *Col3a1^m1Lsmi^* mouse model, respectively.

Given that our measurements demonstrated reduced aortic rupture force of established hAD mouse models in this study (*Fbn1^C1041G^*, *Fbn1*^mgR^, and *Efemp2* SMKO) and elsewhere (*Col3a1^m1Lsmi^*),^12,16^ we evaluated aortic rupture force in three CRISPR/Cas9-engineered mouse models of hAD candidate genes as well. In humans, these three genes (*LTBP1*, *MFAP4*, and *TIMP1*) are expressed at higher levels in arterial tissue than in any other tissue studied in GTEx (gtexportal.org/home). LTBP1 interacts with ECM proteins, such as MFS-associated FBN1, and regulates transforming growth factor (TGF)-β,^47^ which plays an important role in the pathogenesis of hADs.^48^ MFAP4, an ECM protein found in elastic fibers, interacts with FBN1, FBN2, and type I collagen as well as actively promotes tropoelastin self-assembly by binding tropoelastin.^49^ TIMP1 is a strong inhibitor of many MMPs, thereby preventing excessive ECM degradation.^50^ Knock-out mouse models of *Ltbp1*, *Mfap4*, and *Timp1* showed cardiovascular phenotypes,^51–56^ indicating that mutations in these genes could be causative for hAD.

In contrast to established hAD mouse models (i.e., mice with mutation in the genes *Fbn1*, *Efemp2*, or *Col3a1*), however, our read-out system revealed no significantly reduced aortic rupture force for any genotypes of the three novel knock-in mouse models *Ltbp1^G1132S^*, *Mfap4^D42A^*, and *Timp1^I120T^*at the age of 1 year (Figure 5). This indicates and clarifies the lack of aortic significance of the human-equivalent sequence variants with unknown pathogenicity knocked into our mouse models. The non-reduced aortic rupture forces of the three knock-in mouse models are consistent with their non-increased aortic stretch and non-reduced survival rate (data not shown). Further studies are needed to assess the clinical significance of the genes *LTBP1*, *MFAP4*, and *TIMP1* in hAD, also considering a disease-modifying (synergistic) effect of these genes as well as mice with a different genetic background.^57^

Taken together, this study demonstrates that the measurement of aortic rupture force can not only explain the aortic phenotype of established mouse hAD models but also clarify the unknown/uncertain aortic significance of candidate genes, variants, and drugs, thereby paving the way for better molecular diagnosis and medical therapies in hAD. Thus, rupture force can be a valuable parameter to consider in future studies of aortic aneurysms and dissections, especially in evaluating the beneficial effects of pharmacological therapies to attenuate the risk of these life-threatening conditions.

## Acknowledgements

We thank Cyagen (Santa Clara, CA, USA) for help with the generation of the three CRISPR/Cas9 knock-in mouse models, Bryana M. Levitan for performing ultrasound measurements of *Fbn1^mgR^*mice as well as Daniela Brkan, Edina Jusufi, Marc T. Schönholzer, Giancarlo Tomio, and Dzenneta Zukovic for help with data analysis and/or figure preparation.

## Sources of Funding

This work was supported by the Isaac Dreyfus-Bernheim Stiftung and the Fondation pour la Recherche et le Traitement Médical (FRTM) as well as by NIH NHLBI (grant number K01 HL149984) and JSPS KAKENHI (grant number JP20H03762).

## Disclosures

None.

## Supplemental Material

Statistical Analyses

Figures S1-S3

Tables S1-S12

Major Resources Table

## Nonstandard Abbreviations and Acronyms

hAD(s): hereditary aortic disease(s)
AGTR1: angiotensin II receptor type 1
*Col3a1*: collagen, type III, alpha-1 (encoding gene in mice)
ECM: extracellular matrix
*Efemp2*: fibulin-4 (Fbln4) (encoding gene in mice)
*Fbn1*: fibrillin-1 (encoding gene in mice)
*Ltbp1*: latent TGF-β binding protein 1 (encoding gene in mice)
*Mfap4*: microfibril associated protein 4 (encoding gene in mice)
MFS: Marfan syndrome
MMP: matrix metalloproteinase
SMKO: smooth-muscle-specific knockout (*Efemp2^loxP/KO^;SM22-Cre^Tg/Tg^*)
TAA(s): thoracic aortic aneurysm(s)
TGF: transforming growth factor
*Timp1*: tissue inhibitor of matrix metalloproteinase 1 (encoding gene in mice)
vEDS: vascular Ehlers-Danlos syndrome
WGS: whole-genome sequencing

## Notes

### Competing Interest Statement

The authors have declared no competing interest.

### Summary of Updates

Text updated to clarify; Supplemental Material updated.

## REFERENCES

1. Lindsay ME, Dietz HC. Lessons on the pathogenesis of aneurysm from heritable conditions. Nature. 2011;473:308–316. doi: 10.1038/nature10145

2. Verstraeten A, Luyckx I, Loeys B. Aetiology and management of hereditary aortopathy. Nat Rev Cardiol. 2017;14:197–208. doi: 10.1038/nrcardio.2016.211

3. Creamer TJ, Bramel EE, MacFarlane EG. Insights on the pathogenesis of aneurysm through the study of hereditary aortopathies. Genes. 2021;12:183. doi: 10.3390/genes12020183

4. Caspar SM, Dubacher N, Kopps AM, Meienberg J, Henggeler C, Matyas G. Clinical sequencing: From raw data to diagnosis with lifetime value. Clin Genet. 2018;93:508–519. doi: 10.1111/cge.13190

5. Renard M, Francis C, Ghosh R, Scott AF, Witmer PD, Adès LC, Andelfinger GU, Arnaud P, Boileau C, Callewaert BL, Guo D, Hanna N, Lindsay ME, Morisaki H, Morisaki T, Pachter N, Robert L, Van Laer L, Dietz HC, Loeys BL, Milewicz DM, De Backer J. Clinical Validity of Genes for Heritable Thoracic Aortic Aneurysm and Dissection. J Am Coll Cardiol. 2018;72:605–615. doi: 10.1016/j.jacc.2018.04.089

6. Caspar SM, Dubacher N, Matyas G. More Genes for Thoracic Aortic Aneurysms and Dissections. J Am Coll Cardiol. 2019;73:528–529. doi: 10.1016/j.jacc.2018.09.094

7. Humphrey JD, Milewicz DM, Tellides G, Schwartz MA. Dysfunctional mechanosensing in aneurysms. Science. 2014;344:477–479. doi: 10.1126/science.1253026

8. Humphrey JD, Schwartz MA, Tellides G, Milewicz DM. Role of mechanotransduction in vascular biology: focus on thoracic aortic aneurysms and dissections. Circ Res. 2015;116:1448–1461. doi: 10.1161/CIRCRESAHA.114.304936

9. Pierce DM, Maier F, Weisbecker H, Viertler C, Verbrugghe P, Famaey N, Fourneau I, Herijgers P, Holzapfel GA. Human thoracic and abdominal aortic aneurysmal tissues: Damage experiments, statistical analysis and constitutive modeling. J Mech Behav Biomed Mater. 2015;41:92–107. doi: 10.1016/j.jmbbm.2014.10.003

10. Chiu P, Lee H-P, Dalal AR, Koyano T, Nguyen M, Connolly AJ, Chaudhuri O, Fischbein MP. Relative strain is a novel predictor of aneurysmal degeneration of the thoracic aorta: An ex vivo mechanical study. JVS Vasc Sci. 2021;2:235–246. doi: 10.1016/j.jvssci.2021.08.003

11. Chittajallu SNSH, Richhariya A, Tse KM, Chinthapenta V. A Review on Damage and Rupture Modelling for Soft Tissues. Bioengineering (Basel). 2022;9:26. doi: 10.3390/bioengineering9010026

12. Dubacher N, Münger J, Gorosabel MC, Crabb J, Ksiazek AA, Caspar SM, Bakker ENTP, van Bavel E, Ziegler U, Carrel T, Steinmann B, Zeisberger S, Meienberg J, Matyas G. Celiprolol but not losartan improves the biomechanical integrity of the aorta in a mouse model of vascular Ehlers-Danlos syndrome. Cardiovasc Res. 2019;16:457–465. doi: 10.21037/acs.2017.11.10

13. Ong K-T, Perdu J, De Backer J, Bozec E, Collignon P, Emmerich J, Fauret A-L, Fiessinger J-N, Germain DP, Georgesco G, Hulot J-S, De Paepe A, Plauchu H, Jeunemaitre X, Laurent S, Boutouyrie P. Effect of celiprolol on prevention of cardiovascular events in vascular Ehlers-Danlos syndrome: a prospective randomised, open, blinded-endpoints trial. Lancet 2010;376:1476–1484. doi: 10.1016/S0140-6736(10)60960-9

14. Frank M, Adham S, Seigle S, Legrand A, Mirault T, Henneton P, Albuisson J, Denarié N, Mazzella J-M, Mousseaux E, Messas E, Boutouyrie P, Jeunemaitre X. Vascular Ehlers-Danlos Syndrome: Long-Term Observational study. J Am Coll Cardiol. 2019;73:1948–1957. doi: 10.1016/j.jacc.2019.01.058

15. Baderkhan H, Wanhainen A, Stenborg A, Stattin E-L, Björck M. Celiprolol Treatment in Patients with Vascular Ehlers-Danlos Syndrome. Eur J Vasc Endovasc Surg. 2021;61:326–331. doi: 10.1016/j.ejvs.2020.10.020

16. Gorosabel MC, Dubacher N, Meienberg J, Matyas G. Vascular Ehlers-Danlos syndrome: can the beneficial effect of celiprolol be extrapolated to bisoprolol? EHJ - Cardiovasc Pharmacother. 2020;6:199–200. doi: 10.1093/ehjcvp/pvz067

17. Judge DP, Biery NJ, Keene DR, Geubtner J, Myers L, Huso DL, Sakai LY, Dietz HC. Evidence for a critical contribution of haploinsufficiency in the complex pathogenesis of Marfan syndrome. J Clin Invest. 2004;114:172–181. doi: 10.1172/JCI20641

18. Pereira L, Lee SY, Gayraud B, Andrikopoulos K, Shapiro SD, Bunton T, Biery NJ, Dietz HC, Sakai LY, Ramirez F. Pathogenetic sequence for aneurysm revealed in mice underexpressing fibrillin-1. Procl Natl Acad Sci U S A. 1999;96:3819–3823. doi: 10.1073/pnas.96.7.3819

19. Bunton TE, Biery NJ, Myers L, Gayraud B, Ramirez F, Dietz HC. Phenotypic Alteration of Vascular Smooth Muscle Cells Precedes Elastolysis in a Mouse Model of Marfan Syndrome. Circ Res. 2001;88:37–43. doi: 10.1161/01.res.88.1.37

20. Saeyeldin A, Zafar MA, Velasquez CA, Ip K, Gryaznov A, Brownstein AJ, Li Y, Rizzo JA, Erben Y, Ziganshin BA, Elefteriades JA. Natural history of aortic root aneurysms in Marfan syndrome. Ann Cardiothorac Surg. 2017;6:625–632. doi: 10.21037/acs.2017.11.10

21. Habashi JP, Judge DP, Holm TM, Cohn RD, Loeys BL, Cooper TK, Myers L, Klein EC, Liu G, Calvi C, Podowski M, Neptune ER, Halushka MK, Bedja D, Gabrielson K, Rifkin DB, Carta L, Ramirez F, Huso DL, Dietz HC. Losartan, an AT1 Antagonist, Prevents Aortic Aneurysm in a Mouse Model of Marfan Syndrome. Science. 2006;312:117–121. doi: 10.1681/ASN.2006050508

22. Habashi JP, Doyle JJ, Holm TM, Aziz H, Schoenhoff F, Bedja D, Chen Y, Modiri AN, Judge DP, Dietz HC. Angiotensin II Type 2 Receptor Signaling Attenuates Aortic Aneurysm in Mice Through ERK Antagonism. Science. 2011;332:361–365. doi: 10.1126/science.1192152

23. Bhatt AB, Buck JS, Zuflacht JP, Milian J, Kadivar S, Gauvreau K, Singh MN, Creager MA. Distinct effects of losartan and atenolol on vascular stiffness in Marfan syndrome. Vasc Med. 2015;20:317–325. doi: 10.1177/1358863X15569868

24. Cook JR, Clayton NP, Carta L, Galatioto J, Chiu E, Smaldone S, Nelson CA, Cheng SH, Wentworth BM, Ramirez F. Dimorphic effects of transforming growth factor-β signaling during aortic aneurysm progression in mice suggest a combinatorial therapy for Marfan syndrome. Arterioscler Thromb Vasc Biol. 2015;35:911–917. doi: 10.1161/ATVBAHA.114.305150

25. Robertson DM, Truong DT, Cox DA, Carmichael HL, Ou Z, Minich LL, Williams RV, Selamet Tierney ES. Pediatric Heart Network Trial of Losartan vs. Atenolol in Children and Young Adults with Marfan Syndrome: Impact on Prescription Practices. Pediatr Cardiol. 2023;44:618–623. doi: 10.1007/s00246-022-02976-z

26. Huang J, Davis EC, Chapman SL, Budatha M, Marmorstein LY, Word RA, Yanagisawa H. Fibulin-4 Deficiency Results in Ascending Aortic Aneurysms: A Potential Link Between Abnormal Smooth Muscle Cell Phenotype and Aneurysm Progression. Circ Res. 2010;106:583–592. doi: 10.1161/CIRCRESAHA.109.207852

27. Hucthagowder V, Sausgruber N, Kim KH, Angle B, Marmorstein LY, Urban Z. Fibulin- 4: A Novel Gene for an Autosomal Recessive Cutis Laxa Syndrome. Am J Hum Genet. 2006;78:1075–1080. doi: 10.1086/504304

28. Renard M, Holm T, Veith R, Callewaert BL, Adès LC, Baspinar O, Pickart A, Dasouki M, Hoyer J, Rauch A, Trapane P, Earing MG, Coucke PJ, Sakai LY, Dietz HC, De Paepe AM, Loeys BL. Altered TGFβ signaling and cardiovascular manifestations in patients with autosomal recessive cutis laxa type I caused by fibulin-4 deficiency. Eur J Hum Genet. 2010;18:895–901. doi: 10.1038/ejhg.2010.45

29. Muthu ML, Tiedemann K, Fradette J, Komarova S, Reinhardt DP. Fibrillin-1 regulates white adipose tissue development, homeostasis, and function. Matrix Biol. 2022;110:106–128. doi: 10.1016/j.matbio.2022.05.002

30. Lee VS, Halabi CM, Hoffman EP, Carmichael N, Leshchiner I, Lian CG, Bierhals AJ, Vuzman D, Brigham Genomic Medicine, Mecham RP, Frank NY, Stitziel NO. Loss of function mutation in *LOX* causes thoracic aortic aneurysm and dissection in humans. Proc Natl Acad Sci U S A. 2016;113:8759–8764. doi: 10.1073/pnas.1601442113

31. Halabi CM, Broekelmann TJ, Lin M, Lee VS, Chu M-L, Mecham RP. Fibulin-4 is essential for maintaining arterial wall integrity in conduit but not muscular arteries. Sci Adv. 2017;3:e1602532. doi: 10.1126/sciadv.1602532

32. Parker SJ, Stotland A, MacFarlane E, Wilson N, Orosco A, Venkatraman V, Madrid K, Gottlieb R, Dietz HC, Van Eyk JE. Proteomics reveals Rictor as a noncanonical TGF-β signaling target during aneurysm progression in Marfan mice. Am J Physiol Heart Circ Physiol. 2018;315:H1112–H1126. doi: 10.1152/ajpheart.00089.2018

33. Bhushan R, Altinbas L, Jäger M, Zaradzki M, Lehmann D, Timmermann B, Clayton NP, Zhu Y, Kallenbach K, Kararigas G, Robinson PN. An integrative systems approach identifies novel candidates in Marfan syndrome-related pathophysiology. J Cell Mol Med. 2019;23:2526–2535. doi: 10.1111/jcmm.14137

34. Bowen CJ, Calderón Giadrosic JF, Burger Z, Rykiel G, Davis EC, Helmers MR, Benke K, Gallo MacFarlane E, Dietz HC. Targetable cellular signaling events mediate vascular pathology in vascular Ehlers-Danlos syndrome. J Clin Invest. 2020;130:686–698. doi: 10.1172/JCI130730

35. Xiong W, Meisinger T, Knispel R, Worth JM, Baxter BT. MMP-2 Regulates Erk1/2 Phosphorylation and Aortic Dilatation in Marfan Syndrome. Circ Res. 2012;110:e92–e101. doi: 10.1161/CIRCRESAHA.112.268268

36. Sheppard MB, Chen JZ, Rateri DL, Moorleghen JJ, Weiland M, Daugherty A. Renin-Angiotensin System Inhibitors Do Not Improve Survival in Fibrillin-1 Hypomorphic Mice with Established Aortic Aneurysm. J Clin Trans Sci. 2019;3:112–113. doi: 10.1017/cts.2019.258

37. Chen JZ, Rateri D, Sheppard M, Daugherty A. Abstract 20315: Sexual Dimorphism in a Marfan Syndrome Mouse Model. Circulation. 2017;136:A20315.

38. Chen JZ, Sawada H, Moorleghen JJ, Weiland M, Daugherty A, Sheppard MB. Aortic strain correlates with elastin fragmentation in fibrillin-1 hypomorphic mice. Circ Rep. 2019;1:199–205. doi: 10.1253/circrep.CR-18-0012

39. Détaint D, Faivre L, Collod-Beroud G, Child AH, Loeys BL, Binquet C, Gautier E, Arbustini E, Mayer K, Arslan-Kirchner M, Stheneur C, Halliday D, Beroud C, Bonithon-Kopp C, Claustres M, Plauchu H, Robinson PN, Kiotsekoglou A, De Backer J, Adès L, Francke U, De Paepe A, Boileau C, Jondeau G. Cardiovascular manifestations in men and women carrying a FBN1 mutation. Eur Heart J. 2010;31:2223–2229. doi: 10.1093/eurheartj/ehq258

40. Holmes KW, Maslen CL, Kindem M, Kroner BL, Song HK, Ravekes W, Dietz HC, Weinsaft JW, Roman MJ, Devereux RB, Pyeritz RE, Bavaria J, Milewski K, Milewicz D, LeMaire SA, Hendershot T, Eagle KA, Tolunay HE, Desvigne-Nickens P, Silberbach M, for the GenTAC Registry Consortium. GenTAC registry report: Gender differences among individuals with genetically triggered thoracic aortic aneurysm and dissection. Am J Med Genet. 2013;161:779–786. doi: 10.1002/ajmg.a.35836

41. Renard M, Muiño-Mosquera L, Manalo EC, Tufa S, Carlson EJ, Keene DR, De Backer J, Sakai LY. Sex, pregnancy and aortic disease in Marfan syndrome. PLoS ONE. 2017;12:e0181166. doi: 10.1371/journal.pone.0181166

42. Horiguchi M, Inoue T, Ohbayashi T, Hirai M, Noda K, Marmorstein LY, Yabe D, Takagi K, Akama TO, Kita T, Kimura T, Nakamura T. Fibulin-4 conducts proper elastogenesis via interaction with cross-linking enzyme lysyl oxidase. Proc Natl Acad Sci U S A. 2009;106:19029–19034. doi: 10.1073/pnas.0908268106

43. Papke CL, Yanagisawa H. Fibulin-4 and fibulin-5 in elastogenesis and beyond: Insights from mouse and human studies. Matrix Biol. 2014;37:142–149. doi: 10.1016/j.matbio.2014.02.004

44. Papke CL, Tsunezumi J, Ringuette L-J, Nagaoka H, Terajima M, Yamashiro Y, Urquhart G, Yamauchi M, Davis EC, Yanagisawa H. Loss of fibulin-4 disrupts collagen synthesis and maturation: implications for pathology resulting from *EFEMP2* mutations. Hum Mol Genet. 2015;24:5867–5879. doi: 10.1093/hmg/ddv308

45. Ruddy JM, Jones JA, Spinale FG, Ikonomidis JS. Regional heterogeneity within the aorta: Relevance to aneurysm disease. J Thorac Cardiovasc Surg. 2008;136:1123–1130. doi: 10.1016/j.jtcvs.2008.06.027

46. Jana S, Hu M, Shen M, Kassiri Z. Extracellular matrix, regional heterogeneity of the aorta, and aortic aneurysm. Exp Mol Med. 2019;51:1–15. doi: 10.1038/s12276-019-0286-3

47. Morisaki T, Morisaki H. Genetics of hereditary large vessel diseases. J Hum Genet. 2016;61:21–26. doi: 10.1038/jhg.2015.119

48. Neptune ER, Frischmeyer PA, Arking DE, Myers L, Bunton TE, Gayraud B, Ramirez F, Sakai LY, Dietz HC. Dysregulation of TGF-β activation contributes to pathogenesis in Marfan syndrome. Nat Genet. 2003;33:407–411. doi: 10.1038/ng1116

49. Pilecki B, Holm AT, Schlosser A, Moeller JB, Wohl AP, Zuk AV, Heumüller SE, Wallis R, Moestrup SK, Sengle G, Holmskov U, Sorensen GL. Characterization of Microfibrillar-associated Protein 4 (MFAP4) as a Tropoelastin- and Fibrillin-binding Protein Involved in Elastic Fiber Formation. J Biol Chem. 2016;291:1103–1114. doi: 10.1074/jbc.M115.681775

50. Brew K, Nagase H. The tissue inhibitors of metalloproteinases (TIMPs): An ancient family with structural and functional diversity. Biochim Biophys Acta. 2010;1803:55–71. doi: 10.1016/j.bbamcr.2010.01.003

51. Roten L, Nemoto S, Simsic J, Coker ML, Rao V, Baicu S, Defreyte G, Soloway PJ, Zile MR, Spinale FG. Effects of Gene Deletion of the Tissue Inhibitor of the Matrix Metalloproteinase-type 1 (TIMP-1) on Left Ventricular Geometry and Function in Mice. J Mol Cell Cardiol. 2000;32:109–120. doi: 10.1006/jmcc.1999.1052

52. Eskandari MK, Vijungco JD, Flores A, Borensztajn J, Shively V, Pearce WH. Enhanced abdominal aortic aneurysm in TIMP-1-deficient mice. J Surg Res. 2005;123:289–293. doi: 10.1016/j.jss.2004.07.247

53. Todorovic V, Frendewey D, Gutstein DE, Chen Y, Freyer L, Finnegan E, Liu F, Murphy A, Valenzuela D, Yancopoulos G, Rifkin DB. Long form of latent TGF-β binding protein 1 (Ltbp1L) is essential for cardiac outflow tract septation and remodeling. Development. 2007;134:3723–3732. doi: 10.1242/dev.008599

54. Todorovic V, Finnegan E, Freyer L, Zilberberg L, Ota M, Rifkin DB. Long form of latent TGF-β binding protein 1 (Ltbp1L) regulates cardiac valve development. Dev Dyn. 2011;240:176–187. doi: 10.1002/dvdy.22521

55. Schlosser A, Pilecki B, Hemstra LE, Kejling K, Kristmannsdottir GB, Wulf-Johansson H, Moeller JB, Füchtbauer E-M, Nielsen O, Kirketerp-Møller K, Dubey LK, Hansen PBL, Stubbe J, Wrede C, Hegermann J, Ochs M, Rathkolb B, Schrewe A, Bekeredjian R, Wolf E, Gailus-Durner V, Fuchs H, Hrabě de Angelis M, Lindholt JS, Holmskov U, Sorensen GL. MFAP4 Promotes Vascular Smooth Muscle Migration, Proliferation and Accelerates Neointima Formation. Arterioscler Thromb Vasc Biol. 2016;36:122–133. doi: 10.1161/ATVBAHA.115.306672

56. Arpino V, Brock M, Gill SE. The role of TIMPs in regulation of extracellular matrix proteolysis. Matrix Biol. 2015;44–46:247-254. doi: 10.1016/j.matbio.2015.03.005

57. Ritskes-Hoitinga M, Leenaars C, Beumer W, Coenen-de Roo T, Stafleu F, Meijboom FLB. Improving Translation by Identifying Evidence for More Human-Relevant Preclinical Strategies. Animals. 2020;10:1170. doi: 10.3390/ani10071170

