## Supplemental Material for "Novel Insights into the Aortic Mechanical Properties of Mice Modeling Hereditary Aortic Diseases"

**Short title:** Aortic Rupture Force in Mice Modeling Aortic Diseases

**Correspondence to:** Gabor Matyas, PhD, Center for Cardiovascular Genetics and Gene Diagnostics, Swiss Foundation for People with Rare Diseases, Wagistrasse 25, 8952 Schlieren-Zurich, Switzerland., Phone +41-43-433-86-86

### SUPPLEMENTAL METHODS

#### Statistical analyses

For arithmetic means, 95% confidence intervals (CIs) were calculated using two-tailed critical values of *t*-distribution. To be more cautious/conservative in assessing significant differences between groups of samples compared in the text, we only considered means with non- or slightly (<2% of means) overlapping 95% CIs. *P* values (rounded to 4 digits) for differences between two means were calculated by using Student's *t*-test (equal sample variances) or Welch's t-test (unequal sample variances), assuming normal distribution and independence of samples. Sample variances were tested using *F*-test. Statistical computations were performed using VassarStats (vassarstats.net) and, to verify data entry, confirmed by independent analysis using R or GraphPad Prism 9 (GraphPad Software, La Jolla, CA, USA). Moreover, to test all pairwise comparisons among means Tukey's procedure was also applied and considered but not included into the results (as less cautious/conservative in assessing significant differences).

For proportions, 95% CIs were calculated using VassarStats with continuity correction (http://vassarstats.net/prop1.html).

### SUPPLEMENTAL TABLES

**Table S1. Primers for genotyping of the CRISPR/Cas9 knock-in mouse models**

| Mouse Model | Forward Primer | Reverse Primer |
|---|---|---|
| *Ltbp1*$^{emG1132S}$ | 5′-TGCACATCGCTTAATGAAAGGTC-3′ | 5′-CAAAGATTGCCTCAGACTGTACCA-3′ |
| *Mfap4*$^{emD42A}$ | 5′-AGCCCCCATTCACAGACAC-3′ | 5′-GTCCCCATGAAGATGAAAAATC-3′ |
| *Timp1*$^{emI120T}$ | 5′-CATGGAAAGCCTCTGTGGATATGC-3′ | 5′-GTCACAAAGGAAAGTTAGACAGACAGCG-3′ |

**Table S2. Relative aortic rupture forces of the *Fbn1*$^{C1041G}$ mouse model**

| Genotype | S1 | S2 | S3 |
|---|---|---|---|
| **Wild-type** | 100.0 (95.4-104.6) %<br>n=10 | 63.7 (60.9-66.5) %<br>n=8 | 66.8 (60.3-73.4) %<br>n=9 |
| ***Fbn1*$^{+/C1041G}$** | 76.6 (71.3-82.0) %<br>n=9 | 54.3 (49.2-59.4) %<br>n=10 | 51.5 (45.0-58.1) %<br>n=10 |

Relative aortic rupture force (%) of three aortic segments (S1-S3) of 10-week-old wild-type (*Fbn1*$^{+/+}$) and *Fbn1*$^{+/C1041G}$ mice (Figure 1A). Data are means with 95% confidence intervals in parentheses. The sample size (n) is indicated and damaged segments and outliers (>2× standard deviation) were excluded.

**Table S3. Stretch at 5 mN of the *Fbn1*$^{C1041G}$ mouse model**

| Genotype | S1 | S2 | S3 |
|---|---|---|---|
| **Wild-type** | 995 (936-1054) µm<br>n=10 | 724 (675-772) µm<br>n=8 | 697 (643-750) µm<br>n=9 |
| ***Fbn1*$^{+/C1041G}$** | 1066 (977-1155) µm<br>n=9 | 802 (746-859) µm<br>n=10 | 797 (701-894) µm<br>n=10 |

Stretch (in µm) at 5 mN (load-free aortic diameter) of three aortic segments (S1-S3) of 10-week-old wild-type (*Fbn1*$^{+/+}$) and *Fbn1*$^{+/C1041G}$ mice (Figure 1B). Data are means with 95% confidence intervals in parentheses. The sample size (n) is indicated (damaged segments and outliers, i.e. >2× standard deviation, were excluded).

**Table S4. Stretch at maximum aortic rupture force of the *Fbn1*$^{C1041G}$ mouse model**

| Genotype | S1 | S2 | S3 |
|---|---|---|---|
| **Wild-type** | 2928 (2763-3093) µm<br>n=10 | 2510 (2335-2685) µm<br>n=8 | 2303 (2060-2545) µm<br>n=9 |
| ***Fbn1*$^{+/C1041G}$** | 2725 (2547-2904) µm<br>n=9 | 2417 (2219-2615) µm<br>n=10 | 2212 (2026-2397) µm<br>n=10 |

Stretches at maximum force (µm) of three aortic segments (S1-S3) of 10-week-old wild-type (*Fbn1*$^{+/+}$) and *Fbn1*$^{+/C1041G}$ mice (Figure 1B). Data are means with 95% confidence intervals in parentheses. The sample size (n) is indicated (damaged segments and outliers, i.e. >2× standard deviation, were excluded).

**Table S5. Relative aortic rupture forces of the *Fbn1*$^{mgR}$ mouse model after 4-week-long losartan treatment**

| Genotype (treatment) | S1 | S2 | S3 |
|---|---|---|---|
| **Wild-type (untreated)** | 100.0 (94.0-106.0) % n=10 | 55.7 (51.6-59.8) % n=10 | 57.6 (51.0-64.1) % n=10 |
| **Wild-type (losartan)** | 91.5 (87.2-95.8) % n=9 | 48.9 (42.9-54.9) % n=9 | 49.9 (43.2-56.6) % n=9 |
| ***Fbn1*$^{mgR/mgR}$ (untreated)** | 76.0 (70.1-81.9) % n=13 | 46.6 (42.7-50.4) % n=13 | 47.6 (43.2-51.9) % n=13 |
| ***Fbn1*$^{mgR/mgR}$ (losartan)** | 73.6 (67.8-79.4) % n=11 | 40.4 (36.2-44.6) % n=11 | 47.8 (44.1-51.5) % n=11 |

Relative aortic rupture forces (%) of three aortic segments (S1-S3) of untreated wild-type (*Fbn1*$^{+/+}$) and *Fbn1*$^{mgR/mgR}$ control as well as *Fbn1*$^{+/+}$ and *Fbn1*$^{mgR/mgR}$ mice treated for 4 weeks with losartan (Figure 3A). Data are means with 95% confidence intervals in parentheses. The sample size (n) is indicated and damaged segments and outliers (>2× standard deviation) were excluded.

**Table S6. Stretch at 5 mN of the aortic segment S1 in the *Fbn1*$^{mgR}$ mouse model after 4-week-long losartan treatment**

| | S1 | |
|---|---|---|
| **Genotype (treatment)** | **Females** | **Males** |
| **Wild-type (untreated)** | 907 (875-940) µm n=5 | 998 (867-1128) µm n=5 |
| **Wild-type (losartan)** | 1003 (819-1187) µm n=4 | 990 (890-1091) µm n=5 |
| ***Fbn1*$^{mgR/mgR}$ (untreated)** | 1511 (1234-1789) µm n=7 | 2241 (1627-2855) µm n=6 |
| ***Fbn1*$^{mgR/mgR}$ (losartan)** | 1095 (908-1282) µm n=5 | 1517 (977-2057) µm n=6 |

Stretch (in µm) at 5 mN (load-free aortic diameter) of the aortic segment S1 of untreated *Fbn1*$^{+/+}$ (wild-type) and *Fbn1*$^{mgR/mgR}$ control as well *Fbn1*$^{+/+}$ and *Fbn1*$^{mgR/mgR}$ mice treated for 4 weeks with losartan separated according to sex (Figure 3B). Data are means with 95% confidence intervals in parentheses. The sample size (n) is indicated (damaged segments and outliers, i.e. >2× standard deviation, were excluded).

**Table S7. Stretch at maximum rupture force of the aortic segment S1 in the *Fbn1*$^{mgR}$ mouse model after 4-week-long losartan treatment**

| | S1 | |
|---|---|---|
| **Genotype (treatment)** | **Females** | **Males** |
| **Wild-type (untreated)** | 2418 (2306-2530) µm<br>n=5 | 2705 (2443-2968) µm<br>n=5 |
| **Wild-type (losartan)** | 2631 (2266-2996) µm<br>n=4 | 2696 (2654-2737) µm<br>n=5 |
| ***Fbn1*$^{mgR/mgR}$ (untreated)** | 3184 (2671-3697) µm<br>n=7 | 4206 (3334-5078) µm<br>n=6 |
| ***Fbn1*$^{mgR/mgR}$ (losartan)** | 2492 (2280-2704) µm<br>n=5 | 3386 (2597-4176) µm<br>n=6 |

Stretches at maximum force (µm) of the aortic segment S1 of untreated *Fbn1*$^{+/+}$ (wild-type) and *Fbn1*$^{mgR/mgR}$ control as well as *Fbn1*$^{+/+}$ and *Fbn1*$^{mgR/mgR}$ mice treated for 4 weeks with losartan separated according to sex (Figure 3B). Data are means with 95% confidence intervals in parentheses. The sample size (n) is indicated (damaged segments and outliers, i.e. >2× standard deviation, were excluded).

**Table S8. Relative aortic rupture forces of the *Efemp2* SMKO mouse model**

| Genotype | S1 | S2 | S3 |
|---|---|---|---|
| **Wild-type** | 100.0 (88.7-111.3) %<br>n=10 | 56.0 (51.0-61.1) %<br>n=9 | 55.7 (44.5-66.9) %<br>n=9 |
| ***Efemp2* SMKO** | 49.5 (29.9-69.0) %<br>n=9 | 52.8 (47.1-58.5) %<br>n=9 | 50.8 (38.1-63.4) %<br>n=8 |

Relative aortic rupture forces (%) of three aortic segments (S1-S3) of 8-week-old wild-type (*Efemp2*$^{+/+}$;*SM22-Cre*$^{Tg/Tg}$) and *Efemp2* SMKO mice (Figure 4A). Data are means with 95% confidence intervals in parentheses. The sample size (n) is indicated (damaged segments and outliers, i.e. >2× standard deviation, were excluded). SMKO indicates smooth-muscle-cell-specific knockout (*Efemp2*$^{loxP/KO}$;*SM22-Cre*$^{Tg/Tg}$).

**Table S9. Stretch at 5 mN of the *Efemp2* SMKO mouse model**

| Genotype | S1 | S2 | S3 |
|---|---|---|---|
| **Wild-type** | 1043 (958-1127) µm<br>n=10 | 854 (780-928) µm<br>n=9 | 796 (716-876) µm<br>n=9 |
| ***Efemp2* SMKO** | 2735 (2343-3126) µm<br>n=9 | 955 (713-1197) µm<br>n=9 | 779 (678-880) µm<br>n=8 |

Stretch (in µm) at 5 mN (load-free aortic diameter) of three aortic segments (S1-S3) of 8-week-old wild-type (*Efemp2*$^{+/+}$;*SM22-Cre*$^{Tg/Tg}$) and *Efemp2* SMKO mice (Figure 4B). Data are means with 95% confidence intervals in parentheses. The sample size (n) is indicated (damaged segments and outliers, i.e. >2× standard deviation, were excluded). SMKO indicates smooth-muscle-cell-specific knockout (*Efemp2*$^{loxP/KO}$;*SM22-Cre*$^{Tg/Tg}$).

**Table S10. Stretch at maximum aortic rupture force of the *Efemp2* SMKO mouse model**

| Genotype | S1 | S2 | S3 |
|---|---|---|---|
| **Wild-type** | 2811 (2591-3031) µm n=10 | 2496 (2139-2853) µm n=9 | 2330 (2030-2630) µm n=9 |
| ***Efemp2* SMKO** | 4047 (3571-4522) µm n=9 | 2014 (1620-2407) µm n=9 | 1676 (1513-1840) µm n=8 |

Stretches at maximum force (µm) of three aortic segments (S1-S3) of untreated wild-type ($Efemp2^{+/+}$;$SM22\text{-}Cre^{Tg/Tg}$) and *Efemp2* SMKO mice (Figure 4B). Data are means with 95% confidence intervals in parentheses. The sample size (n) is indicated (damaged segments and outliers, i.e. >2× standard deviation, were excluded). SMKO indicates smooth-muscle-cell-specific knockout ($Efemp2^{loxP/KO}$;$SM22\text{-}Cre^{Tg/Tg}$).

**Table S11. Relative aortic rupture force of candidate gene CRISPR/Cas9 knock-in models**

| Genotype | S1 | S2 | S3 |
|---|---|---|---|
| $Ltbp1^{+/+}$ | 100.0 (94.8-105.2) % n=8 | 58.9 (53.5-64.3) % n=9 | 52.2 (44.9-59.6) % n=8 |
| $Ltbp1^{+/G1132S}$ | 92.1 (87.1-97.0) % n=8 | 60.1 (54.7-65.4) % n=9 | 53.2 (45.9-60.5) % n=8 |
| $Ltbp1^{G1132S/G1132S}$ | 94.4 (91.5-97.3) % n=9 | 56.9 (50.7-63.2) % n=9 | 55.0 (47.6-62.4) % n=9 |
| $Mfap4^{+/+}$ | 100.0 (91.6-108.4) % n=6 | 60.0 (54.4-65.6) % n=9 | 63.8 (54.2-73.4) % n=6 |
| $Mfap4^{+/D42A}$ | 89.0 (84.2-93.8) % n=7 | 62.9 (58.5-67.3) % n=8 | 73.9 (67.2-80.6) % n=8 |
| $Mfap4^{D42A/D42A}$ | 93.8 (85.8-101.7) % n=7 | 71.9 (63.9-79.9) % n=8 | 62.4 (53.9-71.0) % n=6 |
| $Timp1^{+/+}$ | 100.0 (91.0-109.0) % n=6 | 74.9 (66.7-83.2) % n=7 | 65.9 (59.5-72.3) % n=7 |
| $Timp1^{+/I120T}$ | 108.1 (102.3-113.8) % n=6 | 74.3 (67.8-80.8) % n=5 | 66.0 (56.8-75.2) % n=6 |
| $Timp1^{I120T/I120T}$ **and** $Timp1^{0/I120T}$ | 103.9 (100.2-107.6) % n=12 | 71.5 (65.9-77.0) % n=11 | 63.7 (59.9-67.4) % n=9 |

Relative aortic rupture forces (%) of three aortic segments (S1-S3) of 1-year-old wild-type and hetero- as well as hemi- and homozygous candidate gene CRISPR/Cas9 knock-in mice (Figure 5). Data are means with 95% confidence intervals in parentheses. The sample size (n) is indicated (damaged segments and outliers, i.e. >2× standard deviation, were excluded).

**Table S12. Relative aortic rupture force of the *Fbn1*$^{C1041G}$, *Fbn1*$^{mgR}$, and *Efemp2* SMKO mouse models compared to *Col3a1*$^{m1Lsmi}$ mice modeling vEDS**

| Genotype | S1 | S2 | S3 |
|---|---|---|---|
| ***Col3a1***$^{+/+}$ | 100.0 (96.5-103.5) % n=11 | 55.8 (51.8-59.9) % n=13 | 55.4 (49.0-61.8) % n=15 |
| ***Col3a1***$^{+/m1Lsmi}$ | 56.3 (50.2-62.4) % n=12 | 39.4 (35.4-43.4) % n=14 | 38.4 (34.2-42.7) % n=11 |
| **WT** (*Fbn1*$^{C1041G}$, *Fbn1*$^{mgR}$, and SMKO) | 100.0 (96.0-104.0) % n=30 | 58.2 (55.6-60.7) % n=27 | 59.9 (55.4-64.5) % n=28 |

Relative aortic rupture forces (%) of three aortic segments (S1-S3) of the mouse vEDS model *Col3a1*$^{m1Lsmi}$ described previously[12] as well as pooled wild-type (WT) controls of the mouse hAD models *Fbn1*$^{C1041G}$, *Fbn1*$^{mgR}$, and *Efemp2* SMKO (Figure 6). Note that relative aortic rupture force values were calculated for each of the mouse AD models *Fbn1*$^{C1041G}$, *Fbn1*$^{mgR}$, and *Efemp2* SMKO to the respective wild-type control and values of mutant mice are listed in the Tables S2, S5, and S8, respectively. Data are means with 95% confidence intervals in parentheses. The sample size (n) is indicated (damaged segments and outliers, i.e. >2× standard deviation, were excluded).

### SUPPLEMENTAL FIGURES

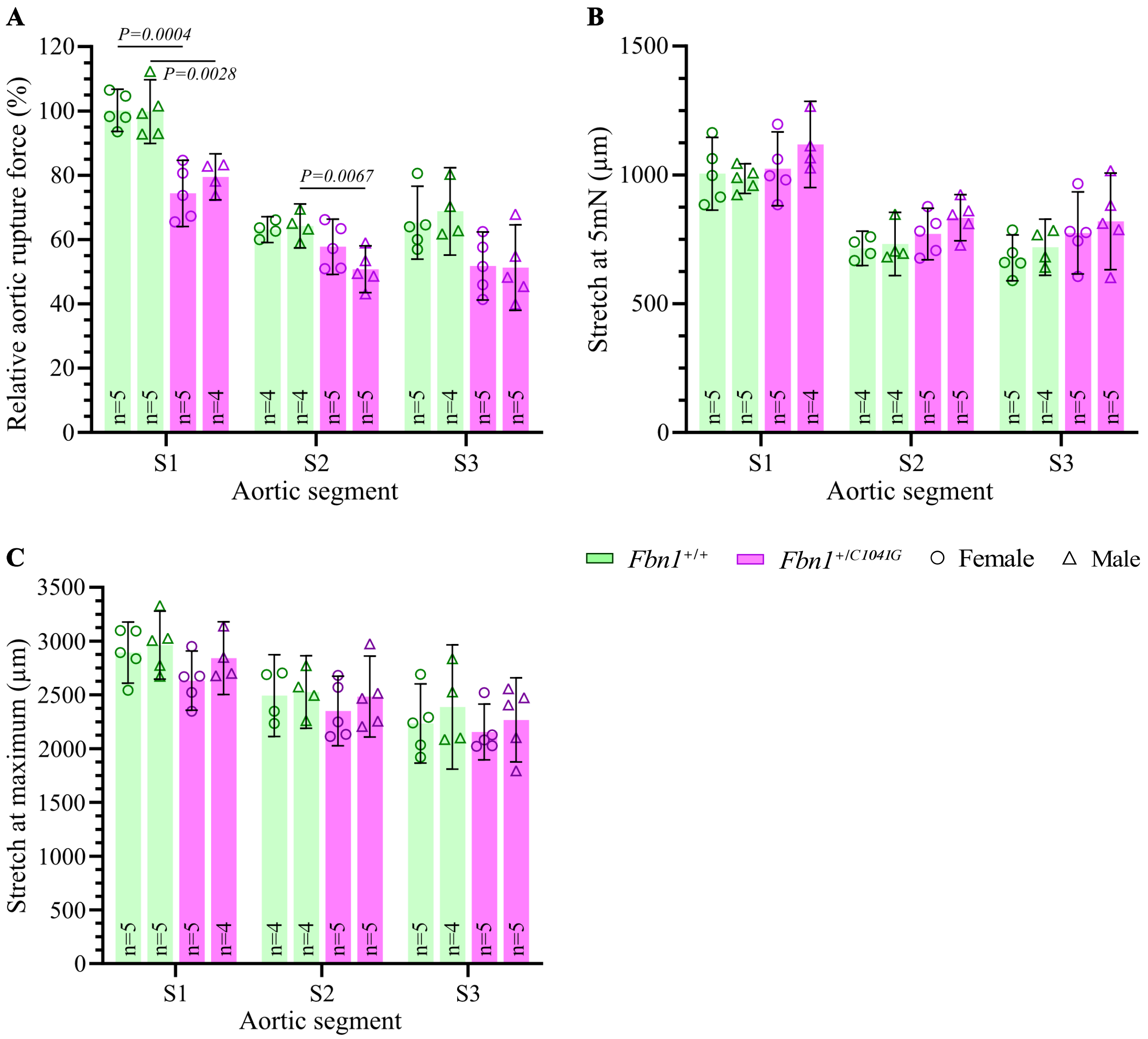


**Figure S1. Aortic parameters of the *Fbn1$^{C1041G}$* mouse model separated according to sex.**
**A**, Relative aortic rupture force (%) of three aortic segments (S1-S3) of 10-week-old *Fbn1$^{+/+}$* and *Fbn1$^{+/C1041G}$* females and males. **B**, Stretch (in µm) of aortic segments at 5mN (load-free aortic diameter) of 10-week-old *Fbn1$^{+/+}$* and *Fbn1$^{+/C1041G}$* females and males. **C**, Stretch (in µm) of aortic segments at maximum force of 10-week-old *Fbn1$^{+/+}$* and *Fbn1$^{+/C1041G}$* mice. Bars represent arithmetic means and error bars indicate 95% confidence intervals. The sample size (n) is displayed (damaged segments and outliers, i.e. >2× standard deviation, were excluded). For mean values with non- or slightly overlapping 95% CIs compared in text, two-tailed *P* values of unpaired t-tests are shown (applies to **A** only).

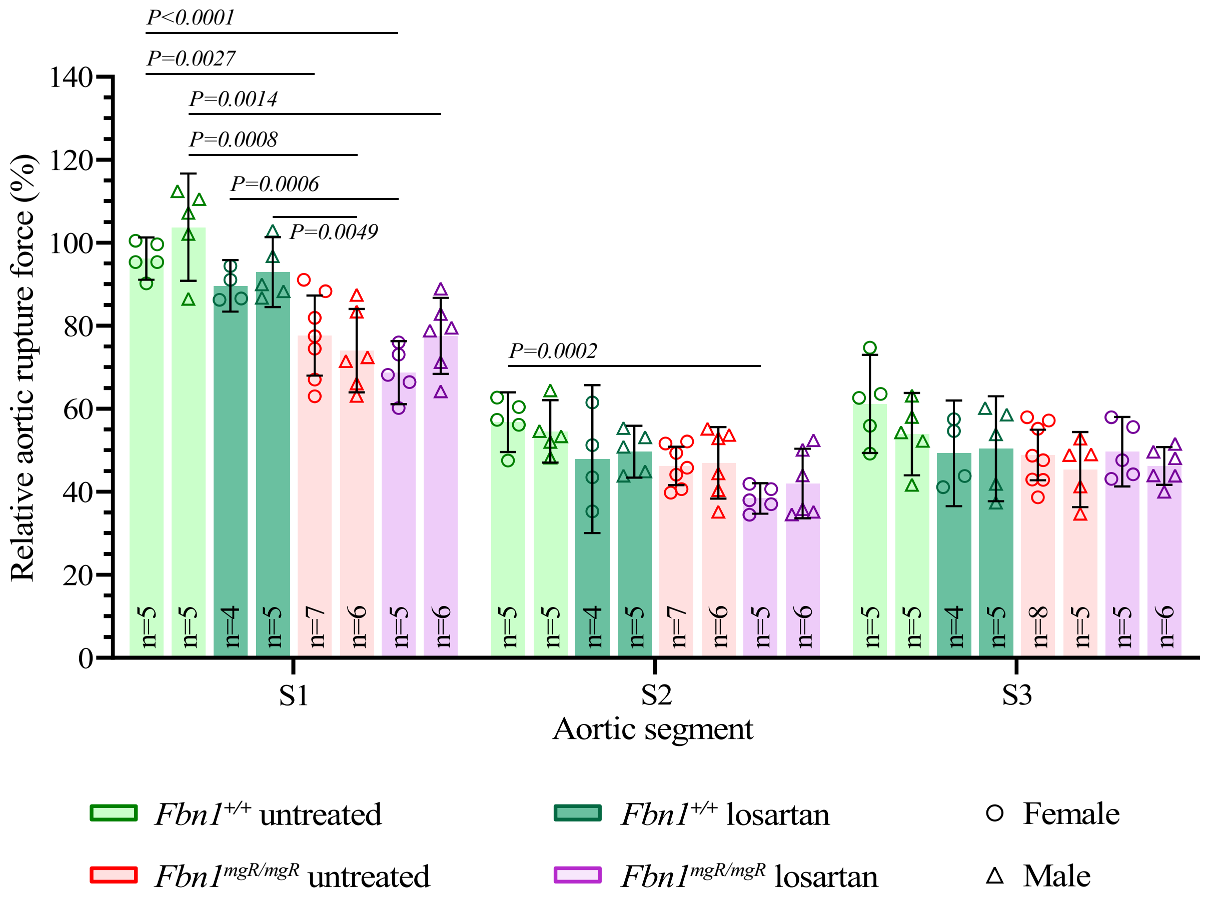


**Figure S2. Aortic rupture force of the *Fbn1^mgR^* mouse model treated with the AGTR1-inhibitor losartan separated according to sex.**

Relative aortic rupture force (%) of three aortic segments (S1-S3) of 8-week-old *Fbn1*$^{+/+}$ and *Fbn1*$^{mgR/mgR}$ females and males after 4-week-long treatment with losartan. Untreated *Fbn1*$^{+/+}$ and *Fbn1*$^{mgR/mgR}$ mice served as control. Bars represent arithmetic means and error bars indicate 95% confidence intervals (95% CI). The sample size (n) is displayed (damaged segments and outliers, i.e. >2× standard deviation, were excluded). For mean values with non- or slightly overlapping 95% CIs compared in text, two-tailed *P* values of unpaired t-tests are shown.

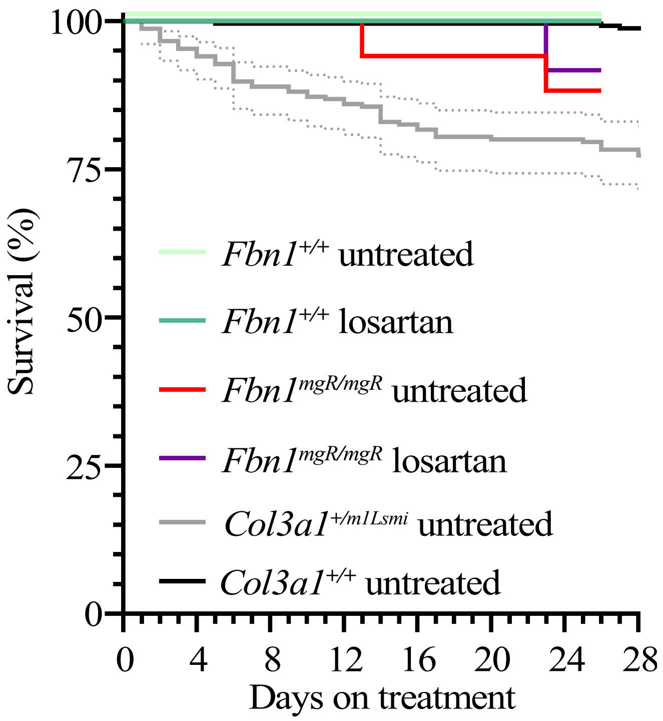


**Figure S3. Survival of the *Fbn1$^{mgR}$* mice modeling MFS and the *Col3a1$^{m1Lsmi}$* mice modeling vEDS**

Kaplan-Meier curve for the survival of untreated *Fbn1$^{+/+}$* (n=10) and *Fbn1$^{mgR/mgR}$* (n=17) mice, *Fbn1$^{+/+}$* (n=10) and *Fbn1$^{mgR/mgR}$* (n=12) mice receiving losartan between the age of 4 and 8 weeks, and untreated *Col3a1$^{+/+}$* (n=241) and *Col3a1$^{+/m1Lsmi}$* (n=235) mice. Note that the survival curves of both losartan-treated *Fbn1$^{+/+}$* mice and untreated *Col3a1$^{+/+}$* mice are partially masked by that of untreated *Fbn1$^{+/+}$* mice. 95% confidence interval (95% CI, dotted line) is shown for *Col3a1$^{+/m1Lsmi}$* mice with survivals differing from 100% (for 95% CI of *Fbn1$^{mgR}$* mice see Figure 2A).

### MAJOR RESOURCES TABLE

Animals

| Species | Vendor or Source | Strain | Genetic Background | Age* |
|---|---|---|---|---|
| Mouse | Jackson Laboratories (Reference ID: IMSR_JAX:012885) | $Fbn1^{C1041G}$ (also known as $Fbn1^{C1039G}$ and $Fbn1^{tm1Hcd}$) | C57BL/6J | 10 weeks |
| | Sheppard lab | $Fbn1^{mgR}$ | C57BL/6J | 8 weeks |
| | Yanagisawa lab | $Efemp2^{loxP/KO}$;$SM22\text{-}Cre^{Tg/Tg}$ (SMKO) | C57BL/6J; 129SvEv | 8 weeks |
| | Cyagen (www.cyagen.com) | $Ltbp1^{emG1132S}$ | C57BL/6NJ | 52 weeks |
| | | $Mfap4^{emD42A}$ | | |
| | | $Timp1^{emI120T}$ | | |

*Age at euthanasia and measurement of the aortic rupture force.
